# *Sphingomonas nivalis* sp. nov., a novel psychrophilic bacteria isolated from a permanent snowfield in Glacier Peaks Wilderness Area, Washington State, USA

**DOI:** 10.1101/2025.06.11.659111

**Authors:** Shawn P. Brown, Amerah Odeh

## Abstract

A novel chemoheterotrophic psychrophilic bacterium, designated as strain I4^T^ was isolated form a permanent snowfield in Washington State, USA. Growth is extremely slow, regardless of temperate (tested between 4°-25° C) but was had marginal optimal growth at 8.5° C. I4^T^ was determined to be able to utilize 102 of the 190 tested carbon sources, 70 or the 95 tested nitrogen sources, 34 or the 59 tested phosphorous sources, and 10 of the 35 tested sulfur sources. I4^T^ was seen to grow from pH 5.0-5.5 but could grow as low as 4.5 pH and as high as 9.5 pH with specific amendments. The genome of I4^T^ was sequenced and it is approximate 4.0 Mb and has a G + C content of 66.2 mol%. Phylogenomic analyses using two independent methods places I4^T^ within the genus *Sphingomonas* with *S. sanguinis* being its closest relative. On the basis of its genomic, physiological, and morphological properties, I4^T^ (=ATCC TSD-357^T^= NCIMB 15472^T^) is proposed as the type strain of a novel species, *Sphingomonas nivalis* sp. nov.

## Introduction

Bacteria have been studied in permanent or semi-permanent snows for over a century (Mclean 1918), and despite the general misconception of snows being devoid of life (Miteva 2008), snows are a habitat for diverse microorganisms (Amato et al. 2007; Brown and Jumpponen 2019; Tucker and Brown 2022). Over the last decade, driven by advances in sequencing technologies, investigations into snow microbial ecology have expanded drastically (Anesio et al. 2017; Brown and Jumpponen 2019; Tucker and Brown 2022; Yakimovich and Quarmby 2022), but culture-based bacteriology continues with several novel snow-born taxa being described recently (Shen et al. 2013; Kojima et al. 2016, 2020). Continued investigations into snow microbiology are important as semi-permanent and perennial snows are a threatened ecosystem due to global climate alterations (Larose et al. 2013; Lutz et al. 2016). Thus, snow can be considered as a vast reservoir of potential novel biodiversity whose environments are threatened (Brown et al. 2015) and this unknown biodiversity may disappear before we understand what may be lost. Bacteria found in snows are diverse, but metabarcoding and other surveys often includes members of the chemoorganotrophic genus *Sphingomonas* (Liu et al. 2006; Chuvochina et al. 2011; Lopatina et al. 2013; Antony et al. 2016).

*Sphingomonas* was proposed by Yabuuchi et al., in 1990 (Yabuuchi et al. 1990) with the type species of *Sphingomonas paucimobilis* (VL34; basonym of *Pseudomonas paucimobilis* (HOLMES et al. 1977)) and the genus has been emended several times (Takeuchi et al. 2001; Yabuuchi et al. 2002; Busse et al. 2003; Chen et al. 2012; Feng et al. 2017). At the time of this writing, there are at least 127 validly published species (Sood et al. 2021). Members of *Sphingomonas* are strictly aerobic, chemoheterotrophic, non-sporulating rod-shaped, Gram-stain-negative taxa. They are known for producing high concentrations of sphingolipids and use ubiquinone-10 (Q-10) as their major respiratory quinone. Many species have pigmented colonies, usually yellow but also orange and red are common but some are translucent and colorless. The present study reports a taxonomic characterization of a novel *Sphingomonas* species, *Sphingomonas nivalis,* isolated from alpine late-season snow.

## Materials and Methods

### Isolation and genome sequencing

The novel species described here was isolated from a perennial snow field (Lyman snowfield; 48° 10’ 17’’ N; 120° 53’ 14’’ W) just downslope from Spider Gap within the Glacier Peaks Wilderness Area in Wenatchee National Forest, Washington State, USA. It was isolated from at north facing slope (30° slope) at an elevation of 2016 ± 12 (m asl) on August 12, 2021. Snow physicochemistry parameters from collection site were measured and the snow had a pH of 8.6 (VIVOSUN digital pH meter; Ontario, CA USA), TDS of 91 ppm, electrical conductivity of 142 of µs/cm (VIVOSUN digital TDS and EC meter; Ontario, CA USA) and ORP value (oxidation-reduction potential) of 138 mV (Gain Express ORP meter, Hong Kong, China).

For isolation, surface snow (top 1 cm) was removed and a 1cm deep surface scraping using a sterile (sterilized with denatured ethanol in the field) aluminum sampling device following (Brown and Jumpponen 2019) and snow was placed into a sterile 50 mL centrifuge tube. Snow was allowed to melt under ambient conditions and 1 mL was plated on a Potato Dextrose Agar (PDA) amended with antibiotics (0.1mL of 10,000U/mL Pen-Strep (ThermoFisher Scientific, Waltham, MA, USA). It is important to note that the goal of this isolation was to generate a snow-borne fungal culture library, which is why PDA plate was amended with antibiotics. The PDA plate was sealed with parafilm and left at ambient temperature in the dark for one day and then incubated at 8.5° C until mixed cultures were visible. Subculturing was conducted on PDA plates and subculturing was continued until a pure culture was obtained. This isolated bacteria (Strain I4) was maintained on PDA at 8.5° C and used for genomic, phenotypic, and physiological interrogation. Type strain (I4^T^) is deposited at the American Type Culture Collection (ATCC; Manassas, VA, USA) under Type Strain Deposit number TDS-357 and at the National Collection of Industrial, Food and Marine Bacteria (NCIMB; Aberdeen, Scotland, UK) under the accession NCIMB 15572).

DNA was extracted using DNeasy Plant Pro extraction kits (Qiagen, Germantown, MD USA), and DNA was quantified using a Qubit 3.0 fluorometer (dsDNA high-sensitivity assay kit Thermo Fisher Scientific, Waltham, MA USA) and 1.0 µg was sequenced on one SpotON flow cell (R9) on the MinION platform (Oxford Nanopore Technologies; Cambridge, UK). Library generation was generated using the ligation kit (SQK-KSK109) and fragments were enriched for long fragments using the long fragment buffer (LFB). Sequence generation was conducted for 72 hours and bases were called using program MiniKnow (Fast Basecalling). The genome was assembled using the long-read assembler Flye (v.2.9; de Bruijn-graph-based)) (Kolmogorov et al. 2019). Total ribosomal regions (16S, 23S, and 5S) were extracted from the whole genome assembly using barrnap (v. 0.9) (Seemann 2013) to aid in quick taxonomic evaluation using a combination of BLASTn and MOLE-BLAST (Altschul et al. 1990) and these initial queries suggested novelty of this taxa and additional investigations were conducted to verify.

## Results and discussion

### Phylogeny and con-generic comparisons

Phylogenetic analyses based on the whole genome assembly was conducted in two different ways. First, we used OrthoFinder (Emms and Kelly 2019) to assess orthrogroups for phylogenetic analyses across the genus *Sphingomonas*. We used every published *Sphingomonas* genome (as of July 2022) and *Hephaestia caeni* (sister genus to *Sphingomonas* (Felföldi et al. 2014)) was used as an outgroup (Table 1). The program fastANI (Jain et al. 2018) was used to determine average nucleotide identify (ANI) between this taxon and other queried taxa. First, predicted protein coding regions of all genomes were identified using Pyrodigal (Larralde 2022) and these were used to identify Othrologs to create individual gene trees and these individual gene trees were used to generate a duplication-loss-coalescence (DCL) resolved gene tree (Figure 1). The genome of this taxon shared the most orthogroups with *S. sanguinis* (2418) and the fewest with *S. sediminicola* (616) and had a median of 1214 shared orthogrops across the phylogeny. This phylogeny clearly demonstrates that this taxon is distinct from, but closely related to *S. sanguinis* and also closely related to *S. paucimobilis* (Figure 1), a finding that the ANI analysis supports (Table 1). Additionally, autoMLST (Alanjary et al. 2019) was used for multi-locus species tree generation, which through MASH distances (Ondov et al. 2016) generate ANI estimates and uses full and partial genomes to select co-occurring genes and builds a maximum-likelihood (ML) coalescent tree, thus more taxa can be included even if whole genome assemblies are not available. In all, autoMLST queried the genome of *S. nivalis* against 11,036 taxa and built a concatenated ML tree a tree using 83 genes and the final tree included 50 terminal taxa nodes (Figure 2, Table S1, Table S2) with *Sphingobium yanoikuyae* (ATTC 51230) as an outgroup (Figure 2). These two methods produced largely congruent relational results with autoMLST placing *S. nivalis* sister to the clade that contains *S. sanguinis, S. paucimobilis*, *S. yabuuchiae,* and several unidentified *Sphingomonas* spp.

**Figure 1.**
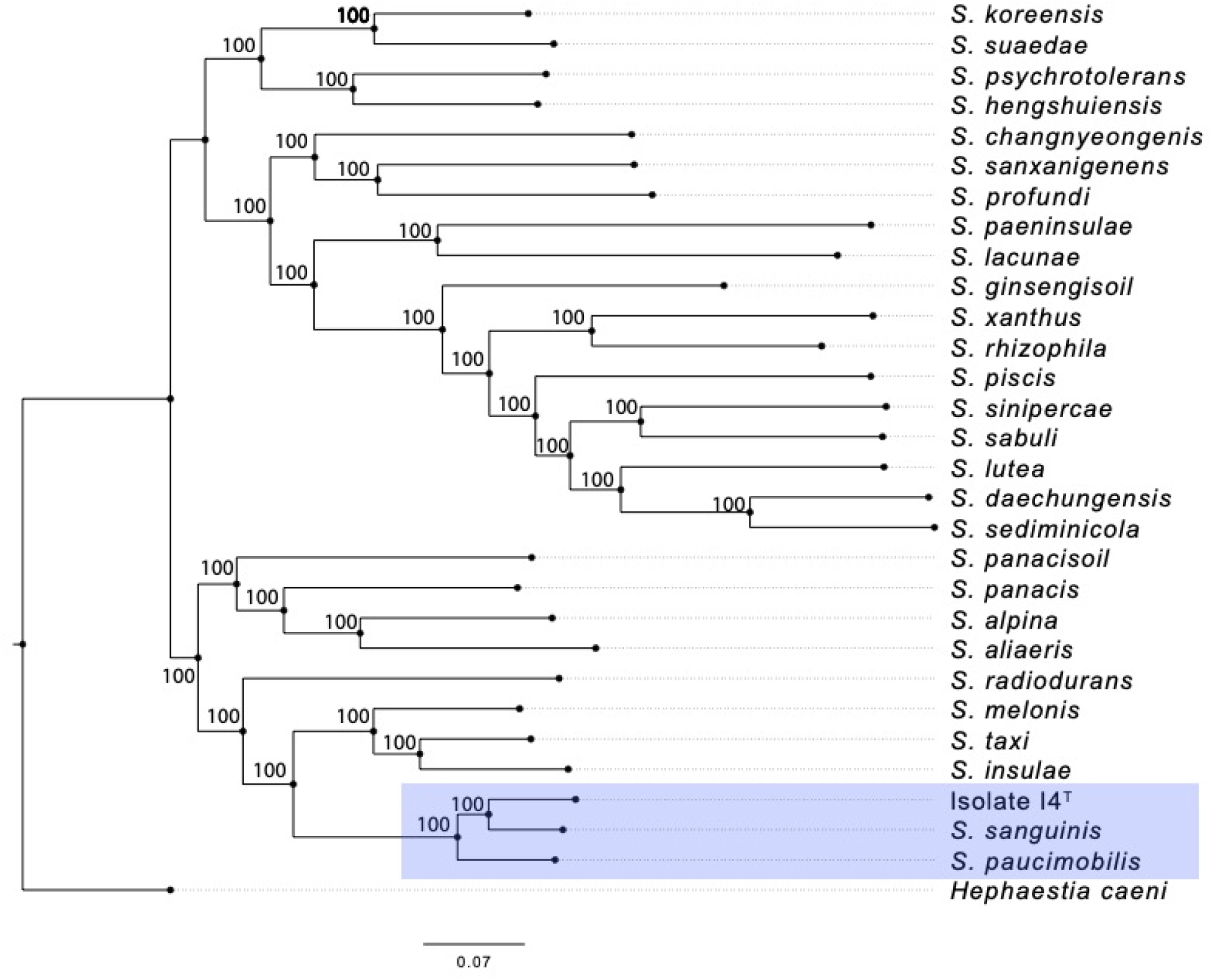
Duplication-loss-coalescence resolved species tree demonstrating complete support (bootstrap) for relationship across *Sphingomonas* indicating that the novel isolate I4^T^ is related to, yet distinct from *S. sanguinis* and *S. paucimobilis* (colored rectangle).

**Figure 2.**
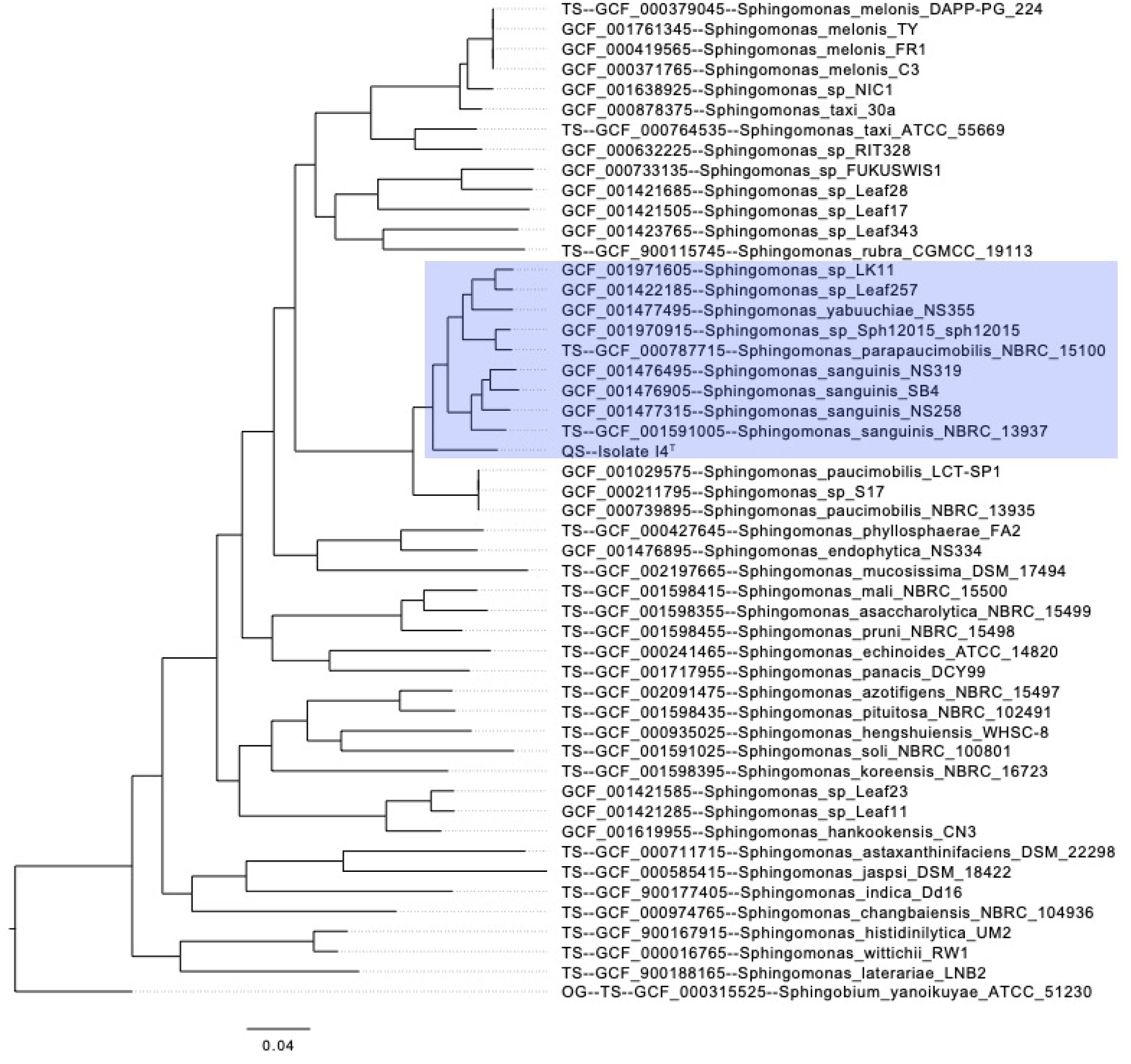
Maximum likelihood tree of the 50 most closely related based on autoMLST analyses demonstrates that strain I4^T^ is sister to *Sphingomonas sanguinis, S. parapaucimobilis, S. yabuuchiae* and several *Sphingomonas* spp. (colored rectangle).

**Table 1.** Genomes used in phylogenomic analyses to generate ortholog based species trees using the program OrthoFinder.

| Species | NCBI Assembly<br>Accession Number | Genome<br>Length<br>(Mb) | GC<br>Content<br>(%) | ANI<br>compared<br>to strain I4<sup>T</sup> |
| --- | --- | --- | --- | --- |
| *Hephaestia caeni* | GCA_003550065.1 | 4.52 | 65.6 | 78.9218 |
| *Sphingomonas sanguinis* | GCA_019297835.1 | 4.14 | 66.1 | 88.1755 |
| *S. paucimobilis* | GCA_016027095.1 | 4.29 | 65.6 | 86.4695 |
| *S. melonis* | GCA_002504265.1 | 3.93 | 67.2 | 80.8455 |
| *S. taxi* | GCA_000764535.1 | 3.86 | 68.0 | 80.3459 |
| *S. insulae* | GCA_010450875.1 | 3.36 | 67.1 | 80.1367 |
| *S. koreensis* | GCA_001922385.1 | 4.51 | 66.1 | 79.1949 |
| *S. panacis* | GCA_001717955.1 | 5.32 | 65.5 | 79.1866 |
| *S. hengshuiensis* | GCA_000935025.1 | 5.19 | 66.7 | 79.1698 |
| *S. suaedae* | GCA_007833215.1 | 4.15 | 65.5 | 79.0825 |
| *S. alpina* | GCA_014490665.1 | 5.10 | 64.0 | 78.9562 |
| *S. aliaeris* | GCA_016743815.1 | 4.26 | 64.4 | 78.9394 |
| *S. changnyeongenensis* | GCA_009913435.1 | 3.01 | 68.3 | 78.9084 |
| *S. panacisoil* | GCA_007859635.1 | 3.40 | 65.1 | 78.8811 |
| *S. radiodurans* | GCA_020866845.1 | 3.67 | 65.7 | 78.7759 |
| *S. sanxanigenens* | GCA_000512205.2 | 6.21 | 66.8 | 78.6703 |
| *S. psychrotolerans* | GCA_002796605.1 | 5.03 | 65.6 | 78.4440 |
| *S. profundus* | GCA_009739515.1 | 4.53 | 69.2 | 78.4179 |
| *S. ginsengisoil* | GCA_009363895.1 | 3.03 | 67.2 | 78.1399 |
| *S. rhizophila* | GCA_014396585.1 | 2.55 | 65.2 | 78.1384 |
| *S. sabuli* | GCA_014352855.1 | 2.51 | 65.9 | 77.8591 |
| *S. xanthus* | GCA_007998985.1 | 2.23 | 63.7 | 77.7910 |
| *S. sinipercae* | GCA_011302055.1 | 2.07 | 65.5 | 77.7719 |
| *S. lutea* | GCA_014396785.1 | 2.42 | 65.3 | 77.7071 |
| *S. piscis* | GCA_011300455.1 | 2.72 | 63.3 | 77.6867 |
| *S. lacunae* | GCA_012979535.1 | 2.97 | 62.0 | 77.6837 |

### Genome features

The genome of *Sphingomonas nivalis* (strain I4^T^) is accessioned at NCBI WGS under the following accessions: BioProject PRJNA940784, BioSample SAMN33578476, and PGAP JARGGQ000000000. Ribosomal sequences are accessioned at NCBI GenBank under the following accessions: 16S (OR130162) and 23S (OR130163). Assembly size is 4,000,803 bp with 129x coverage across 9 contigs with a N50 of 1,081,545, with a G+C content of 66.2%. OrthoFinder (Emms and Kelly 2019) identified 5457 predicted genes and 2782 orthogroups within *S. nivalis*, 956 of which *S. nivalis* possess multiple copies of. There were 105 orthogroups that were not shared with other tested taxa in which include 221 genes. Further, there are 992 predicted gene duplication events in the genome of this taxon, all of which have > 50% support and were all identified as terminal duplications suggesting very little differentiation between duplicated genes. A total of 444 putative xenologs (genes derived from horizontal genes transfer events) were identified in the *S. nivalis* genome, 27 (6.1%) of which are associated with *S. changnyeongensis*, 25 (5.6%) to *S. sanxanigenens* and 25 (5.6%) to *S. profundi*. This is interesting as these three taxa form a monophyletic clade (Figure 1) but are distant to *S. nivalis* so how these are related to HGT events remains unclear.

### Phenotypic characterization, nutrient utilization, and physiology

This taxon is a Gram-negative, yellow-pigmented, very slow-growing bacterium with ovoid cell morphology (Figure 3). Colonies were imaged using Scanning Electron Microscopy (Nova NanoSEM 650; FEI, Hillsboro, OR USA) and additional investigations using an Oxford EDS system (Energy Dispersive X-Ray Spectroscopy) was conducted to examine surface elemental composition of this taxon. Using several SEM images, cell size was measured for 100 cells using ImageJ (Schneider et al. 2012) and cell size is as follows: long axis 1.14 μm ± 0.17 μm (mean ± standard deviation), short axis 0.54 μm ± 0.07 μm, surface area 6.63 μm^2^ ± 1.12 μm^2^, cell volume 1.42 μm^3^ ± 0.36 μm^3^, surface area and cell volume are measured following estimates for prolate spheroid cells following (Ojkic et al. 2019). The measured surface area to volume ratio is 4.77 ± 0.53. In SEM micrographs, no evidence of polar flagella was seen as is the most common with this genus (Sood et al. 2021), so we presume *S. nivalis* has gliding motility as seen in a few other fresh water taxa within this genus (Lee et al. 2016, 2017), but this was not explicitly tested.

**Figure 3.**
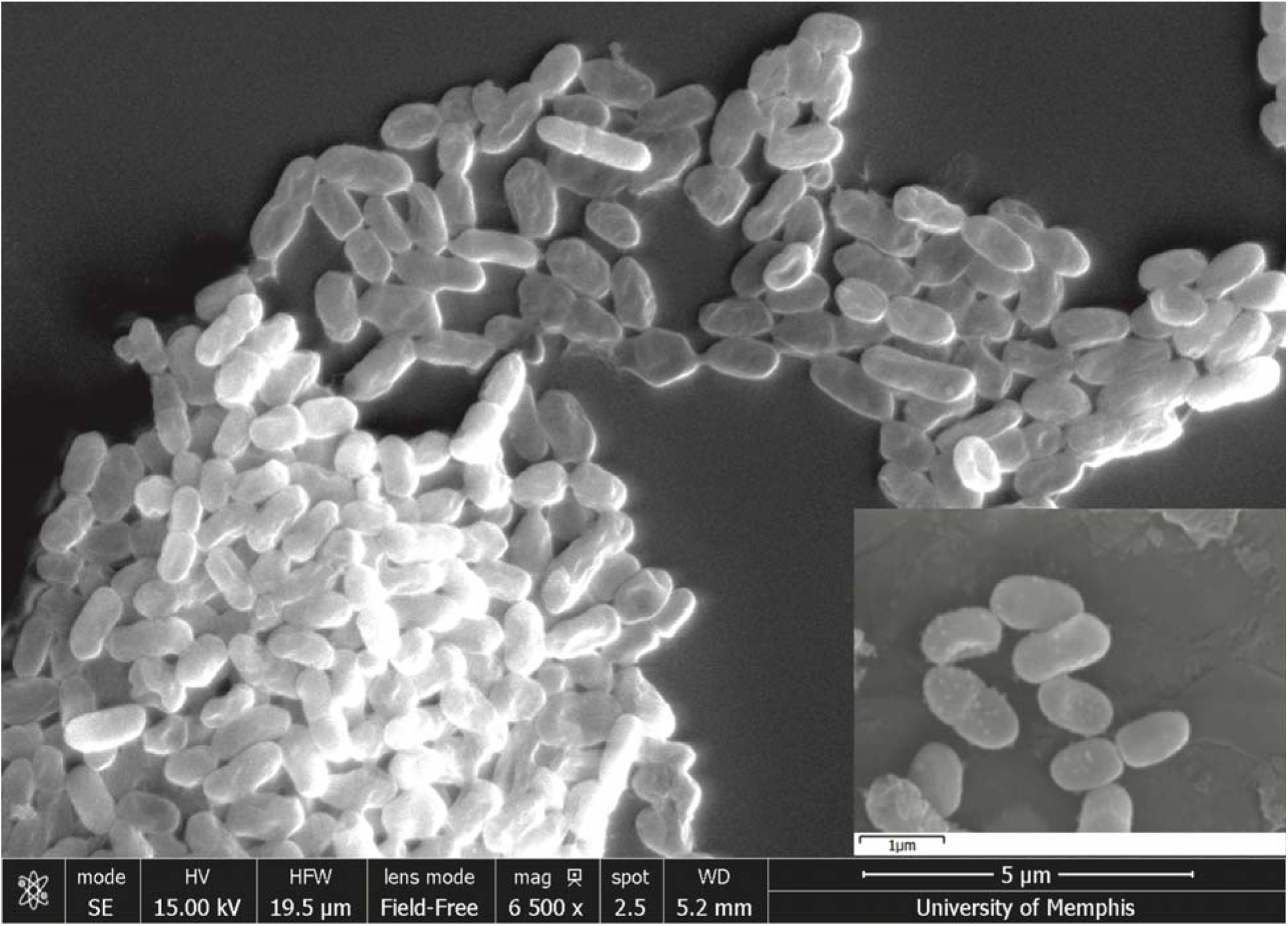
SEM micrographs demonstrating cell morphology of strain I4^T^.

The SEM-EDS analysis suggested that in addition to surface carbon, nitrogen, and oxygen as would be expected, there was enriched Zinc (Zn) and Silicon (Si) associated with the surface of this taxon compared to background scatter. This may indicate that this taxon may preferentially utilize and/or otherwise sequester these elements, or, growth may be enhanced by these elements. Zinc is generally considered toxic to bacteria (Bamberger et al. 1993; McDevitt et al. 2011), even at low concentrations, but many bacteria are zinc tolerant and may even select for zinc sequestration, presumably as a cellular mechanism to immobilize toxic zinc via cell wall modification and bioprecipitation (Choudhury and Srivastava 2001). Zinc biosorption research is primally focused on plant growth promoting bacteria (PGPB) (He et al. 2010; Kour et al. 2019), but general mechanisms of Zn tolerance are poorly understood (Mangold et al. 2013). Silicon utilization by bacteria, which includes biosolubilizing mechanisms (Bist et al. 2020), is a much better understood process and extracellular biosilicification is a common occurrence (Ikeda 2021) which may aid in exopolysaccharide production to aid in protection from harsh environments (Fetsiukh et al. 2021). To test if growth of this taxon is influenced or enhanced by Si or Zn concentrations, we conducted differential growth assays. By plating this taxon in triplicate on PDA media amended with either Zinc Sulfate Heptahydrate (ZnSO_4_) or Sodium Metasilicate Nonahydrate (Na_2_SiO_3_) at concentrations of 0μM, 0.05μM, 0.2μM, 0.5μM, and 1μM of either Zn or Si. Plates were incubated at 8.5° C for 40 days and colony area (mm^2^) were measured using high-resolution photography and ImageJ (Schneider et al. 2012) at 3 d, 7 d, 12 d, 20 d, 25 d, 30 d, 35 d and 40 d. Effects of Zn or Si were tested using a two-way ANOVA (model: days of growth, concentration, and their interaction) and where significant for concentration, Tukey HSD post-hoc tests were conducted to determine which concentrations differed. Both Zn (F_9,120_=43.87, P<0.0001) and Si (F_9,120_=123.53, P<0.0001) had growth that were significantly impacted by our model, with both Zn and Si causing differential growth with concentrations (Zn: F_4,120_=34.05, P<0.0001; Si: F_4,120_=11.85, P<0.0001) and concentration by days of growth interactions (Zn: F_4,120_=5.80, P=0.0003; Si: F_4,120_=2.50, P=0.423). Post hoc tests indicate Zn suppress growth substantially (0μM - 117.98 mm^2^ ± 5.83 mm^2^ ^A^, 0.05μM - 48.36 mm^2^ ± 1.92 mm^2^ ^C^, 0.2μM - 49.12 mm^2^ ± 2.15 mm^2^ ^C^, 0.5μM - 53.55 mm^2^ ± 3.41 mm^2^ ^B,C^, and 1μM - 59.57 mm^2^ ± 3.01 mm^2^ ^B^, Least Square Means ± standard error (mm^2^) and superscript letters represent Tukey HSD connecting letters) and moderate Si suppress growth but high and minimal Si levels were not different from Si-free (0μM - 88.73 mm^2^ ± 2.35 mm^2^ ^A^, 0.05μM - 91.15 mm^2^ ± 2.54 mm^2^ ^A^, 0.2μM – 76.21 mm^2^ ± 2.47 mm^2^ ^B^, 0.5μM – 78.40 mm^2^ ± 2.37 mm^2^ ^B^, and 1μM – 95.94 mm^2^ ± 2.48 mm^2^ ^A^). While growth results were interesting, we find little evidence of enhanced growth with Si and we see reduced growth with Zn.

Nutrition utilization and pH physiological growth capabilities were tested (Table 2) using Biolog Phenotype MicroArrays (Biolog, Hayward, CA). Following standard procedures, capability of this taxon to utilize 190 carbon sources (Biolog plates PM1 and PM2A), 95 nitrogen sources (plate PM3B), 59 phosphorous and 35 sulfur sources (plate PM4A). Further ability to grow across a pH range of 3.5-10 and various pH levels with biological amendments were tested (plate PM10). MicroArray plating was conducted using standard protocols, incubated at 8.5° C for seven days and growth was measured using a SynergyHTX Multi-Model Microplate Reader (BioTek, Winooski, VT) at a wavelength of 590nm. Growth was categorized by comparing OD-590 values to controls and no-growth (-) is where OD-590 < 1.25x control, mild growth (+) is where 1.25x control < OD-590 < 1.5x control, moderate growth (++) is where 1.5x control < OD-590 < 1.75x control, and extreme growth (+++) where OD-590 > 1.75x control. In all, of those tested, this taxon can utilize 102 carbon sources, 70 nitrogen sources, 34 phosphorus sources, and 10 sulfur sources (Table 2). In the pH assays, growth only occurred at pH 5.0 and 5.5 (both +++) but could also grow outside this range with certain amendments (pH 4.5 with Anthranilic acid, +++; pH 4.5 with 5-Hydroxy Tryptophan, +++; pH 9.5 with L-Glutamine, ++; pH 9.5 with L-Tyrosine, +++, pH 9.5 with Phenylethylamine, +++; pH 9.5 with Tyramine, +++). In the pH assay, growth also occurred on X-Caprylate (+++) and X-SO_4_ (+). The other 83 pH levels or amendments were negative for growth.

**Table 2.**
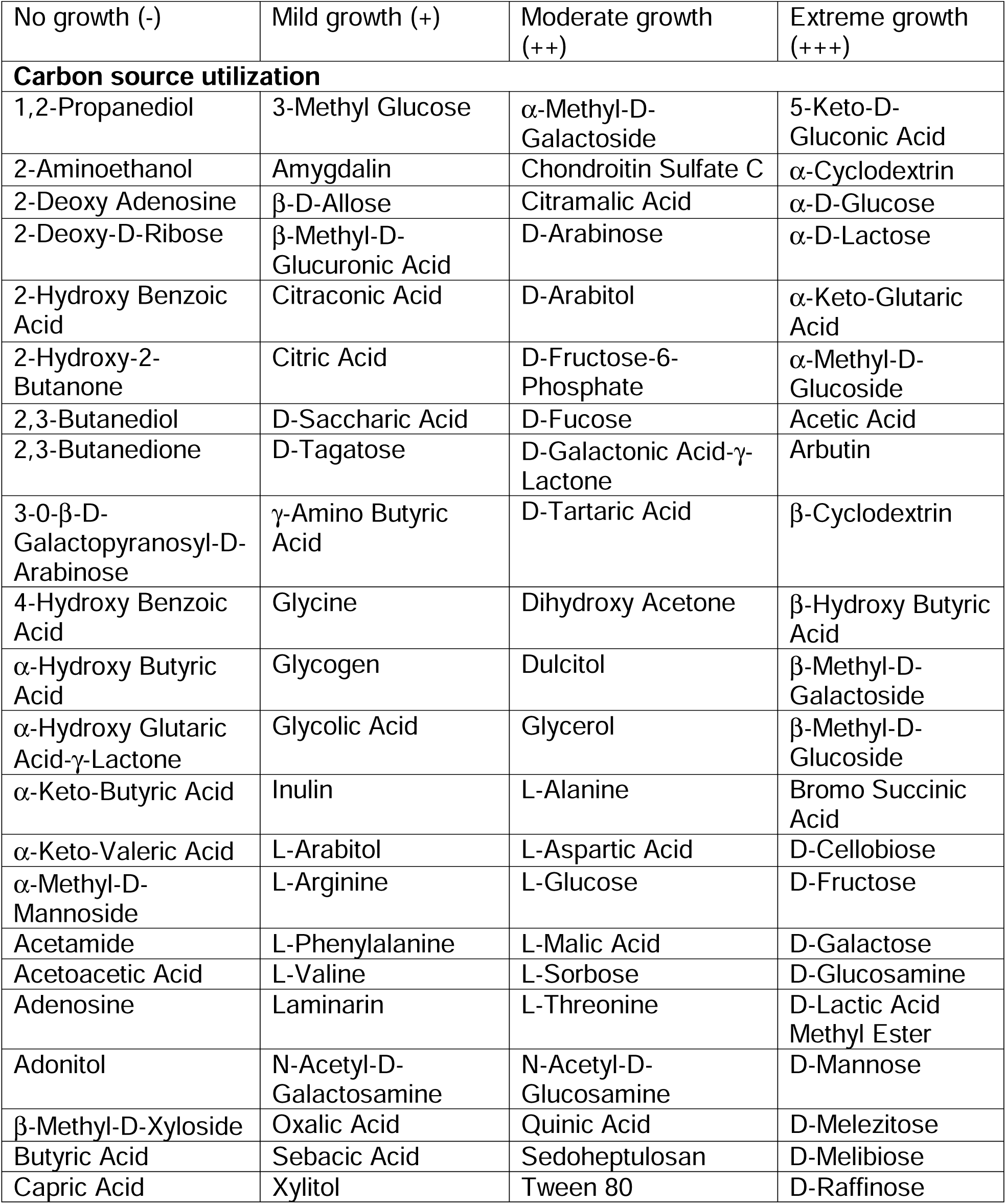
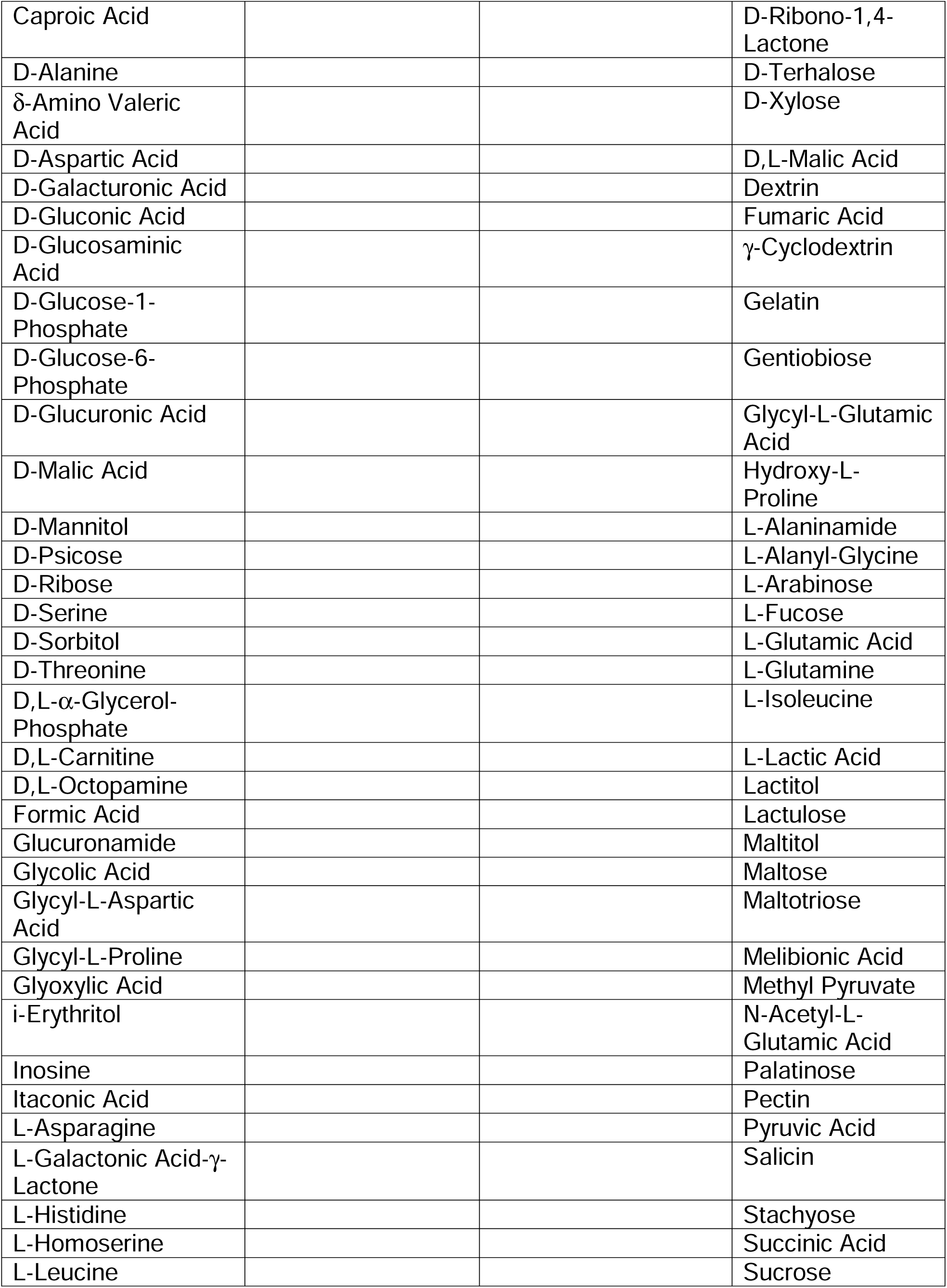
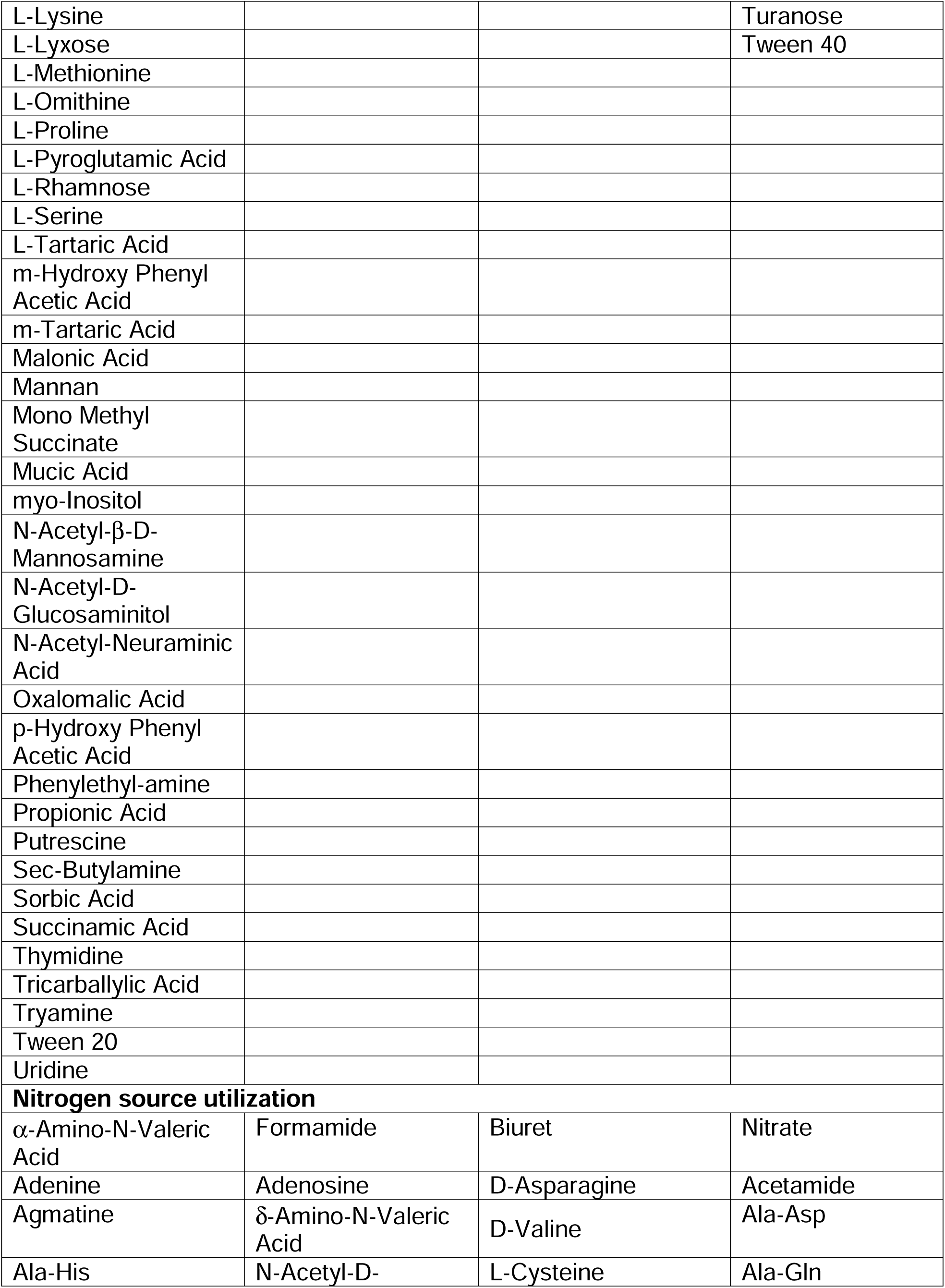
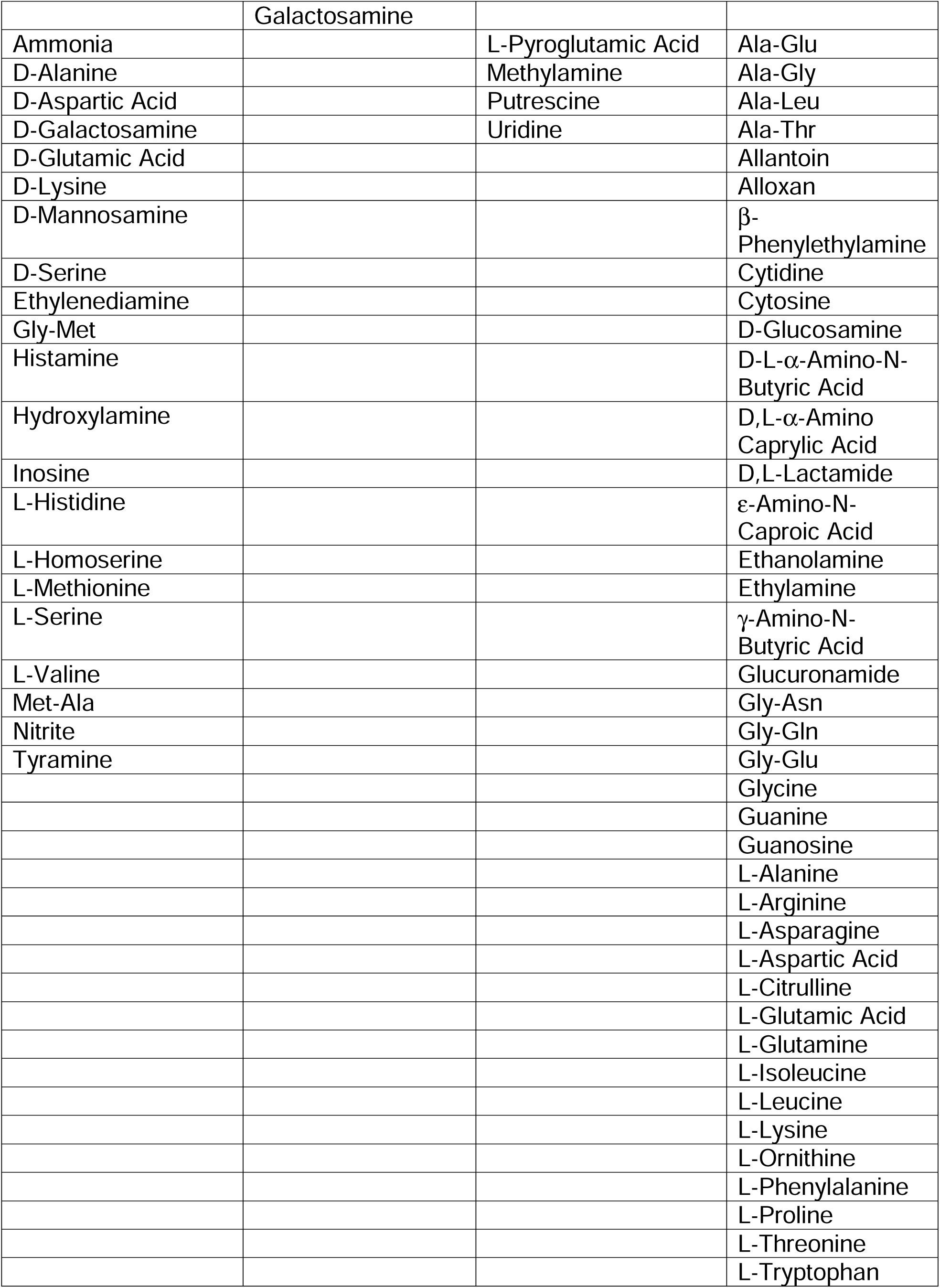
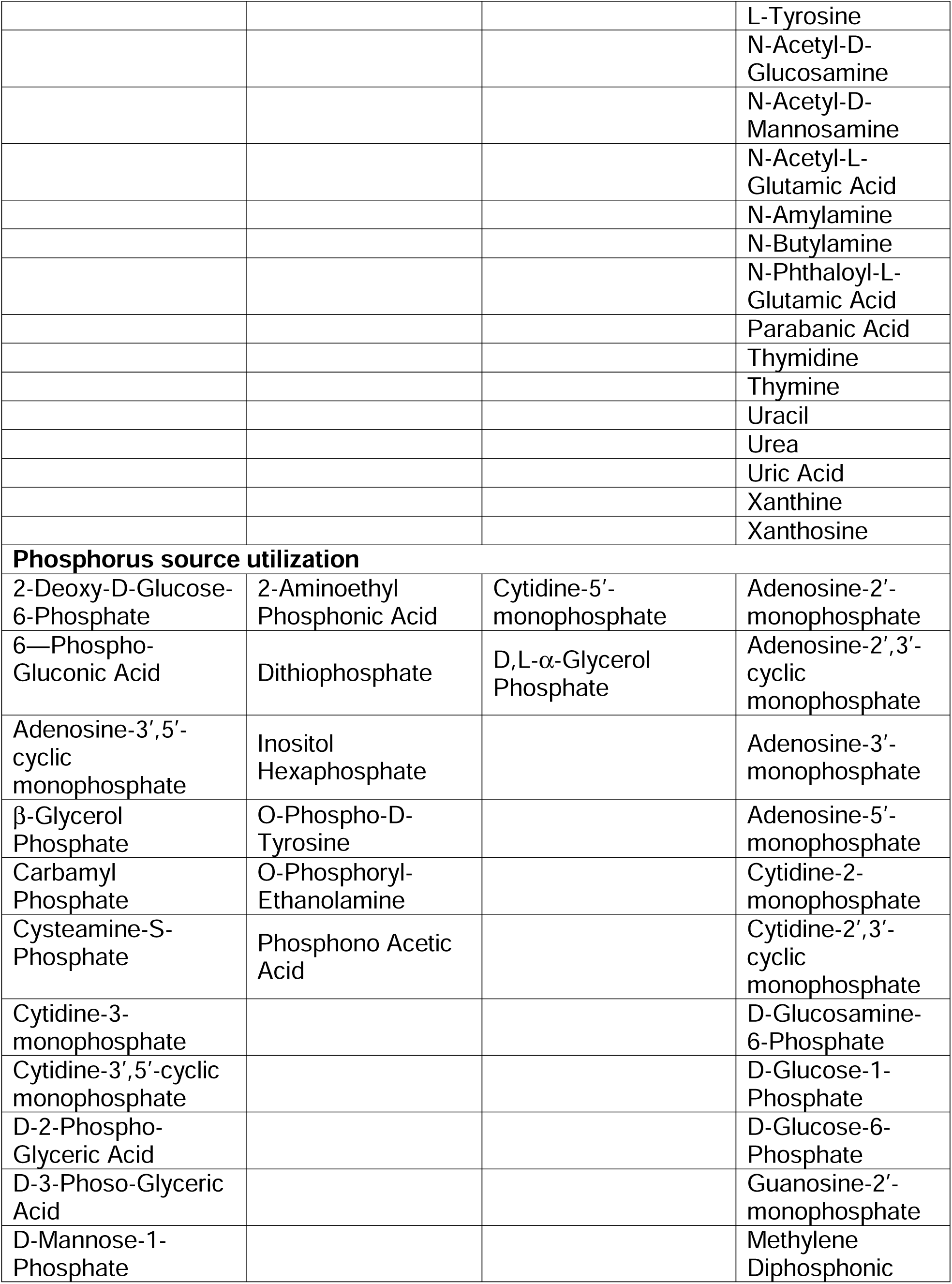
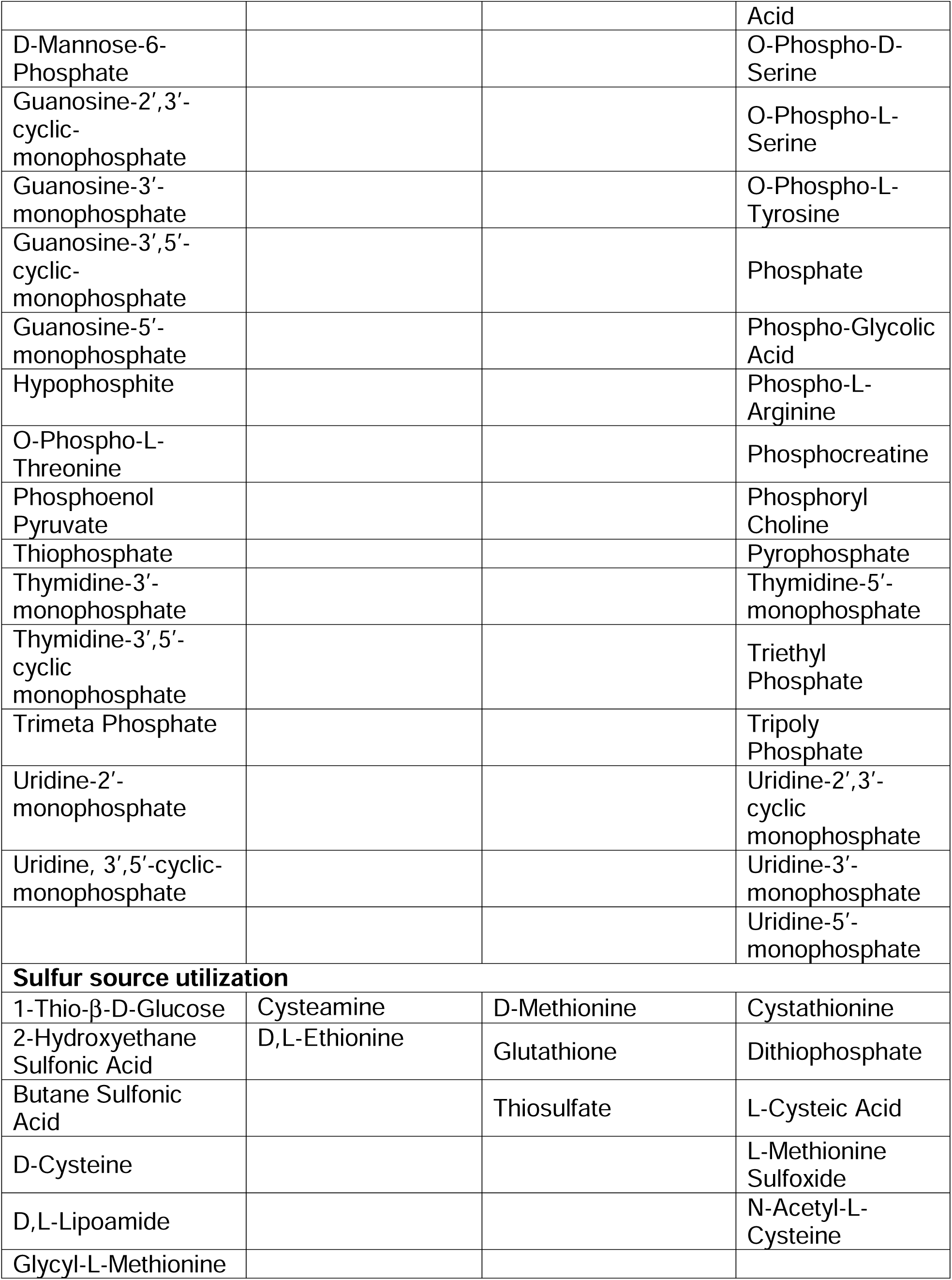
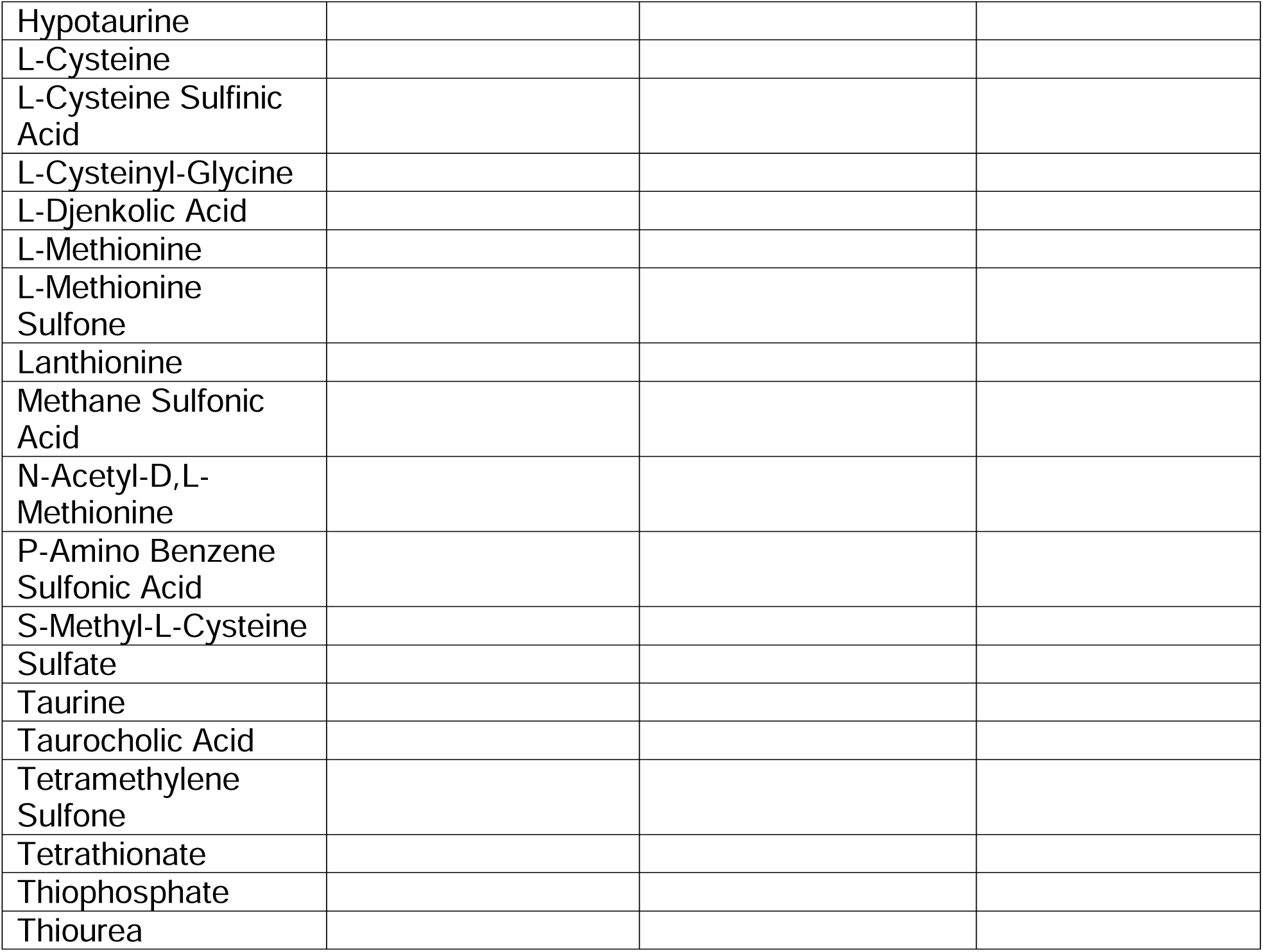
Nutrient utilization and growth capacity properties of strain I4^T^ based on Phenotypic MicroArrays. Included are carbon, nitrogen, phosphorous, and sulfur assays as well as pH. Growth was measured compared to negative controls using OD values at 590nm and how well I4^T^ utilizes elemental source is presented as no growth (-), mild growth (+), moderate growth (++), and extreme growth (+++).

Based on polyphasic investigations into this taxon, including genomic, physiological, and nutrient utilization assays, we propose the description of a novel *Sphingomonas* species, for which we suggest the name *Sphingomonas nivalis* (type strain I4^T^), isolated from perennial snows in the cascade mountains in Washington State, USA.

### Description of sphingomonas nivalis sp. Nov

*Sphingomonas nivalis* (na’val.is L. gen. adj. *nivalis*, snow covered, from which the organism was isolated).

Cells are Gram-stain-negative, aerobic, and rod or prolate spheroid shaped, approximately 1.14 μm in length and 0.54 μm wide. Culture growth on solid PDA and R2A media results in sticky, yellow to yellow-orange irregular, slightly umbonate and circular colonies. Growth at 8.5° C for 7 days on solid media resulted in colony areas of an average of 18 mm^2^. Type strain I4^T^ is able to use as a carbon sources 3-Methyl Glucose, 5-Keto-D-Gluconic Acid, α-Cyclodextrin, α-D-Glucose, α-D-Lactose, α-Keto-Glutaric Acid, α-Methyl-D-Galactoside, α-Methyl-D-Glucoside, Acetic Acid, Amygdalin, Arbutin, β-Cyclodextrin, β-D-Allose, β-Hydroxy Butyric Acid, β-Methyl-D-Galactoside, β-Methyl-D-Glucoside, β-Methyl-D-Glucuronic Acid, Bromo Succinic Acid, Chondroitin Sulfate C, Citraconic Acid, Citramalic Acid, Citric Acid, D-Arabinose, D-Arabitol, D-Cellobiose, D-Fructose, D-Fructose-6-Phosphate, D-Fucose, D-Galactonic Acid-g-Lactone, D-Galactose, D-Glucosamine, D-Lactic Acid Methyl Ester, D-Mannose, D-Melezitose, D-Melibiose, D-Raffinose, D-Ribono-1,4-Lactone, D-Saccharic Acid, D-Tagatose, D-Tartaric Acid, D-Terhalose, D-Xylose, D,L-Malic Acid, Dextrin, Dihydroxy Acetone, Dulcitol, Fumaric Acid, γ-Amino Butyric Acid, γ-Cyclodextrin, Gelatin, Gentiobiose, Glycerol, Glycine, Glycogen, Glycolic Acid, Glycyl-L-Glutamic Acid, Hydroxy-L-Proline, Inulin, L-Alaninamide, L-Alanine, L-Alanyl-Glycine, L-Arabinose, L-Arabitol, L-Arginine, L-Aspartic Acid, L-Fucose, L-Glucose, L-Glutamic Acid, L-Glutamine, L-Isoleucine, L-Lactic Acid, L-Malic Acid, L-Phenylalanine, L-Sorbose, L-Threonine, L-Valine, Lactitol, Lactulose, Laminarin, Maltitol, Maltose, Maltotriose, Melibionic Acid, Methyl Pyruvate, N-Acetyl-D-Galactosamine, N-Acetyl-D-Glucosamine, N-Acetyl-L-Glutamic Acid, Oxalic Acid, Palatinose, Pectin, Pyruvic Acid, Quinic Acid, Salicin, Sebacic Acid, Sedoheptulosan, Stachyose, Succinic Acid, Sucrose, Turanose, Tween 40, Tween 80, and Xylitol. The strain can grow in liquid media at from pH 5.0-5.5. The strain is resistant to penicillin and streptomycin.

The type strain, I4^T^ (=ATCC TSD-357^T^= NCIMB 15472^T^), was isolated from a permanent (semi-permanent) snowfield in Washington State, USA. The DNA G+C content of strain I4^T^ is 66.2 mol%. All sequence data are publicly available at NCBI under the accessions PRJNA940784, OR130163 and OR130162.

## Acknowledgements

The authors would like to thank Ari Jumpponen, Joel Spencer and Vee Brown for their assistance in sample collection. We thank Omar Skalli and Lauren Thompson for their assistance with microscopy. Avery Tucker and Grant Pinlac assisted with culture maintenance and DNA extraction. We would like to acknowledge past and present members of the Chelan, Entiat, Yakama, Okanagan tribes and the confederated tribes of the Colville Reservation, whose ancestral land this taxon was isolated from.

## Ethical approval

This article does not contain any studies with human and/or animal participants.

## Figure and Table Legends

**Table S1** List of genes used for phylogenomic analyses using autoMLST. In all 83 genes were used for these tree generation. Presented are autoMLST gene ID, gene name, gene function, and extend gene name with descriptions.

**Table S2** Queried taxa in autoMLST analyses including 11,036 taxa in order to build ML phylogenetic tree across 83 genes. Presented are reference assembly, name of taxa, MASH distances, ANI values, Genus, Order, and if the queried taxon is a type specimen.

